# A repeatedly evolved mutation in Cryptochrome-1 of subterranean animals alters behavioral and molecular circadian rhythms

**DOI:** 10.1101/2024.09.19.613894

**Authors:** Amruta Swaminathan, Alexander Kenzior, Colin McCoin, Andrew Price, Kyle Weaver, Aurélie Hintermann, NatiCia Morris, Alex C. Keene, Nicolas Rohner

**Affiliations:** Stowers Institute for Medical Research, Kansas City, MO, USA; Department of Cell Biology and Physiology, University of Kansas Medical Center, Kansas City, KS, USA; Department of Biology, Texas A&M University, College Station, TX, USA

## Abstract

The repeated evolution of similar phenotypes in independent lineages often occurs in response to similar environmental pressures, through similar or different molecular pathways. Recently, a repeatedly occurring mutation R263Q in a conserved domain of the protein Cryptochrome-1 (CRY1) was reported in multiple species inhabiting subterranean environments. Cryptochromes regulate circadian rhythms, and glucose and lipid metabolism. Subterranean species show changes to their circadian rhythm and metabolic pathways, making it likely that this mutation in CRY1 contributes to adaptive phenotypic changes. To identify the functional consequences of the CRY1 R263Q mutation, we generated a mouse model homozygous for this mutation. Indirect calorimetry experiments revealed delayed energy expenditure, locomotor activity and feeding patterns of mutant mice in the dark phase, but no further metabolic phenotypes – unlike a full loss of function of CRY1. Gene expression analyses showed altered expression of several canonical circadian genes in the livers of the mutant mice, fortifying the notion that CRY1 R263Q impacts metabolism. Our data provide the first characterization of a novel mutation that has repeatedly evolved in subterranean environments, supporting the idea that shared environmental constraints can drive the evolution of similar phenotypes through similar genetic changes.

## INTRODUCTION

Repeated evolution is the recurrence of similar phenotypic traits in multiple lineages. Often, these lineages inhabit similar environments and may face similar selection pressures to develop these traits (Bolnick et al., 2018; Losos, 2011). The same developmental or physiological pathway can be perturbed at different points to produce the same gross phenotype and therefore genetic changes underlying repeated phenotypes are not necessarily the same in different lineages. However, instances of repeated evolution are also observed at the molecular level. Moran et al. (Moran et al., 2022) reported a mutation in the gene *Cryptochrome-1* (*Cry1*) that results in the substitution of an arginine (R) residue at position 263 in the CRY1 protein with a glutamine (Q) residue in a highly conserved domain (**Figure 1A**). Moreover, this mutation has arisen independently at least 5 times during animal evolution, in lineages that inhabit cave or subterranean environments (**Figure 1B**). Given that this specific mutation has repeatedly occurred in a highly conserved gene across species sharing similar environmental pressures, we hypothesize that it does not result simply in a loss of function, but rather alters a specific function of the CRY1 protein. We refer to this mutation henceforth as the CRY1 R263Q mutation, or simply R263Q.

**Figure 1.**
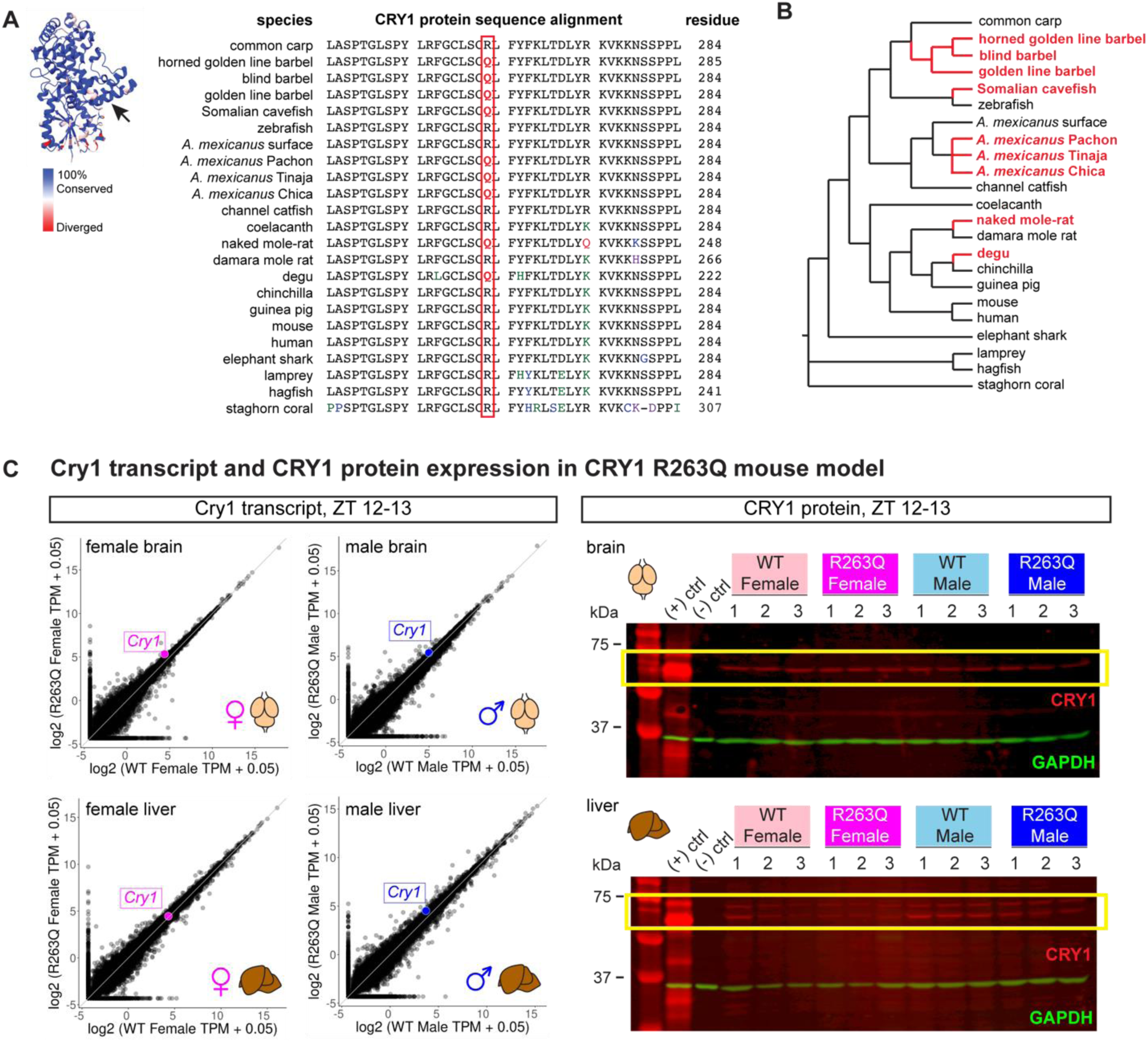
A mouse model to study the repeatedly evolved mutation in Cryptochrome-1 orthologs of subterranean dwellers. 1A. Partial amino acid sequence alignment of CRY1 orthologs from animal lineages including cnidarians, jawless fish, teleosts and mammals. The species is indicated on the left and the residue numbers on the right. The red rectangle marks the site of the R263Q mutation, with the Q highlighted in boldface red font. Alignment redrawn from (Moran et al., 2022). Colored residues indicate amino acid changes. Inset on the left is the crystal structure of mouse Cryptochrome-1 or CRY1 photolyase homology region (PDB: 7D0M). Colors indicate degree of amino acid conservation of mouse CRY1 with human CRY1, zebrafish CRY1a and *Xenopus* CRY1, with blue indicating 100% conservation and red indicating divergence. The arrow marks the position of the CRY1 R263Q mutation in the crystal structure. Structure visualized using ChimeraX 1.5rc202211120143. 1B. Animal phylogeny showing the occurrence of the CRY1 R263Q mutation. The branches in red are lineages in which the mutation has been detected. Phylogeny redrawn from (Moran et al., 2022). 1C. Verifying transcript and protein level expression of the mutant CRY1 R263Q locus. Left: plots of log2 values of mean transcripts per million (TPM) for total mRNA sequenced from CRY1 R263Q mutant versus total mRNA sequenced from CRY1 wild-type male and female brain and liver tissues at zeitgeber time 12-13. Each point represents a single gene. Each combination of animal sex and source tissue is plotted separately. Top left, brain tissue from females; top right, brain tissue from males; bottom left, liver tissue from females; bottom right, liver tissue from males; for brain tissue, *n* = 2 individuals each for wild-type and R263Q mutant for each sex; for liver tissue, *n* = 3 individuals each for wild-type and R263Q mutant for each sex. For each plot, points falling on the grey line (*y* = *x*) represent genes which have equal median TPM values between R263Q mutant individuals and wild-type individuals for a specific sex and tissue. The *Cry1* TPM value (R263Q mutant versus wild-type) for each case is shown as a larger colored point on the plot (magenta for the female plots and blue for the male plots). Right: Western blots showing CRY1 protein expression in the brain (top) and liver (bottom) tissues from wild-type and CRY1 R263Q mice, male and female. The positive control lanes are lysates prepared from U-2 OS CRY1/2 KO human cell lines transfected with an expression vector for mouse CRY1 (pSG5 backbone containing coding sequence for mouse CRY1 with FLAG, His and Myc epitope tags; pMC1SG5 vector as generated and described by (Ye et al., 2011)), and the negative control lanes are lysates prepared from untransfected U-2 OS CRY1/2 KO human cell lines. CRY1 signal is shown in red (inside the yellow rectangles), GAPDH signal is shown in green as a loading control.

CRY1 is one of the major proteins regulating circadian rhythms (Griffin et al., 1999; Kume et al., 1999; Laothamatas et al., 2023; Takahashi, 2017; van der Horst et al., 1999), which are intrinsically generated 24-hour rhythms in the gene expression, physiology, metabolism and behavior of an organism, and are ubiquitous across the tree of life (Dunlap, 1999). Additionally, CRY1 also regulates glucose homeostasis, and lipid and steroid metabolic processes ((Jang et al., 2017; Koike et al., 2012; Lamia et al., 2011; Zhang et al., 2010)). The animals that naturally carry the CRY1 R263Q mutation inhabit caves and underground burrows and are reported to have altered their circadian rhythms and metabolism (Aspiras et al., 2015; Avivi et al., 2001; Avivi et al., 2002, 2004; Cavallari et al., 2011; Ceinos et al., 2018; Di Rosa et al., 2024; Ghosh et al., 2021; Mack et al., 2021; Riccio and Goldman, 2000; Riddle et al., 2018). The finding that the CRY1 R263Q mutation has evolved repeatedly in similar environments suggests a critical role for this mutation in such organisms. To explore this hypothesis, we have generated a mouse line carrying the CRY1 R263Q mutation. We found that CRY1 R263Q mice show delayed energy expenditure, locomotor activity and feeding patterns in the dark phase, but not in the light phase. Gene expression analyses revealed altered expression of major circadian and metabolic genes in the livers of the mutant mice. Taken together, our data suggest that the CRY1 R263Q mutation dysregulates the circadian clock at the transcriptional as well as behavioral levels and may thus aid in adaptation to subterranean life. In a broader context, our findings support the notion that repeated evolution of phenotypes does not necessarily proceed by divergent genetic changes in independent lineages, and it is possible for shared environmental pressures to select for the same genetic change across independent lineages to drive the repeated evolution of a trait.

## MATERIALS AND METHODS

### Mouse strains

CRISPR-Cas9 technology was used for engineering C57BL6/J mice. Potential guide RNA target sites were designed using CCTop (Stemmer et al., 2015). The target site was selected by evaluating the predicted on-target efficiency score and the off-target potential (Labuhn et al., 2017) in addition to the proximity of the double-stranded break to the desired mutation site. To generate the R263Q codon change, a single stranded DNA olignonucleotide (ssODN) donor was designed containing ∼50 nucleotides of homology from the double-stranded break site. The ssODN contained mutations for the R263Q change as well as silent codon changes to disrupt guide RNA binding following the repair event. The ssODN was ordered as an Alt-R HDR Donor Oligo from Integrated DNA Technologies (IDT). The selected guide RNA was ordered as an Alt-R CRISPR-Cas9 sgRNA from IDT. **Figure S1A** provides a graphic summary of the CRISPR design. 3–4-week-old C57BL6/J females were super-ovulated using pregnant mare serum gonadotropin (PMSG) given at noon, followed by human chorionic gonadotropin (HCG) 46 hours later at 10:00 am, and then paired with C57BL6/J stud males overnight. All females with copulation plugs the next morning were kept and used to harvest embryos. Harvested embryos were placed in potassium-supplemented simplex optimized medium (KSOM) and incubated until electroporation was performed.

A ribonucleoprotein (RNP) complex was formed by incubating sgRNA at 6 μM final concentration with IDT Cas9 HiFi v3 protein at 1.2 μM final concentration at room temperature for 10 minutes. The ssODN was added at 200 ng/μL final concentration to the RNP. 84 fertilized embryos were selected and electroporated with the RNP complex using the Nepa Gene electroporator. After electroporation, embryos were transferred into recipient females. These females carried the embryos to term and gave birth to pups which were screened for the expected mutations by lysing an ear clip and amplifying the specific genomic location. A second round of amplification was performed to incorporate sample-specific dual barcodes. All amplicons were pooled and size-selected using ProNex Size-Selective Purification System (Promega). Cleaned pools were quantified on a Qubit Fluorimeter and then run on a Bioanalyzer (Agilent) to check sizing and purity. Purified pools were run on an Illumina MiSeq 2×250 flow cell. The resulting sequence data were demultiplexed, and read pairs were joined. On-target indel frequency and expected mutations were analyzed using CRIS.py (Connelly and Pruett-Miller, 2019).

Founder males were chosen to be paired with C57BL6/J wild-type females to create the F1 generation. The pups resulting from these matings were weaned and sequenced as described above for the founders. Any positive pups with confirmed germline transmission were kept as F1s. When these mice reached breeding age (6-8 weeks) they were set up with new C57BL6/J wild-type mice to produce the F2 generation that contained heterozygotes. Heterozygotes were crossed with each other to produce the F3 generation with litters containing wild-type mice and mice homozygous for the introduced mutation. Since large numbers of animals were required for metabolic experiments, control mice were derived by crossing wild-type mice and the mutant mice were derived by crossing mutant mice. For experiments performed at the Stowers Institute for Medical Research (SIMR), mice were housed in a 14:10 light/dark cycle (light on at 5:45 am and off at 7:45 pm) with food (Teklad 2020X, Inotiv) and water available *ad libitum*. For experiments performed at the University of Kansas Medical Center (KUMC), mice were housed in a 12:12 light/dark cycle (lights on at 6:00 am and off at 6:00 pm) with food (Teklad 8604, Inotiv) and water available *ad libitum* during quarantine periods and experiments. All experiments were performed in accordance with the guidelines of the Institute Animal Care and Use Committees (IACUC) of SIMR and KUMC.

### PCR genotyping

DNA was extracted from ear biopsies using the Qiagen DNeasy Blood and Tissue Kit (Qiagen, 69504) following the manufacturer’s instructions. The site of the CRY1 R263Q mutation was amplified for 35 cycles using a forward primer sequence 5’AGGCCTGGGTGGCAAACTTTG3’ and a reverse primer sequence 5’CGCCACAGGAGTTGCCCATAAAG3’. The primers were designed to amplify a 300 bp fragment flanking the mutation site and spanning the entire ssODN sequence to ensure that the amplification is from the *Cry1* locus and not from elsewhere in the genome. The denaturation, annealing and extension steps were carried out for 30, 30 and 60 seconds respectively, at 95 °C, 60 °C and 72 °C respectively. PCR amplicons were Sanger sequenced to identify wild-type and R263Q mutant samples. **Figure S1B** shows example Sanger sequencing chromatograms comparing the site of mutation in a wild-type individual with a CRY1 R263Q homozygote.

### Tissue lysis and western blotting

Mice housed at SIMR and aged between 3-4 months were euthanized with CO_2_ followed by cervical dislocation. Livers were harvested from each mouse, washed in 1X PBS, and fragments of the largest lobe of each liver were flash frozen in liquid nitrogen. Tissue harvesting was done between ZT12 and ZT13 (onset of the dark phase). For total protein extracts, liver fragments were thawed on ice and then homogenized in 20 μL of radioimmunoprecipitation assay (RIPA) buffer (89900, ThermoFisher Scientific) per mg of tissue, using a Type A dounce homogenizer on ice. Protein concentrations were determined using the BCA assay (23227, Pierce) following the manufacturer’s instructions. Samples were incubated at 70 °C for 10 minutes with 4X Laemmli buffer and then loaded on a 10% polyacrylamide/SDS gel for electrophoresis. The separated proteins were transferred to a 0.45 μm PVDF membrane (IVPH00010, Merck) at a constant voltage of 25 V for 2 hours. Post-transfer, the membrane was washed thrice with TBS-Tw (1X TBS with 0.1% v/v Tween-20) and blocked for 1 hour with a blocking buffer (5% w/v instant nonfat dry milk in TBS-Tw). Primary antibodies were diluted in blocking buffer and incubated with the membrane for 2 hours. Following primary antibody incubation, the membrane was washed thrice with TBS-Tw and then incubated with secondary antibodies diluted in the blocking buffer for 1 hour, then washed thrice. Imaging was carried out on the LiCOR Odyssey CLx system. All steps in the western blotting protocol after transfer were carried out at room temperature. The primary and secondary antibodies and the dilutions used can be found in Table 1.

**Table 1.** Antibodies used in western blotting.

| Catalog # | Antibody | Host/Isotype | Conjugate | Dilution |
| --- | --- | --- | --- | --- |
| PM081 | Anti-CRY1 (mouse) pAb | Guinea pig/IgG | Unconjugated | 1:1,000 |
| 60004-1-Ig | GAPDH monoclonal antibody | Mouse/IgG2 | Unconjugated | 1:10,000 |
| ab1791 | Anti-histone H3 polyclonal antibody | Rabbit/IgG | Unconjugated | 1:10,000 |
| 926-68077 | IRDye® 680RD Donkey anti-Guinea Pig IgG Secondary Antibody | Donkey/IgG | IRDye® 680RD | 1:4,000 |
| 926-32212 | IRDye® 800CW Donkey anti-Mouse IgG Secondary Antibody | Donkey/IgG | IRDye® 800CW | 1:10,000 |
| 926-68073 | IRDye® 680RD Donkey anti-Rabbit IgG Secondary Antibody | Donkey/IgG | IRDye® 680RD | 1:10,000 |

### Indirect calorimetry

Mice aged 3-4 months were transferred from SIMR to the KUMC Metabolic Core and placed under quarantine for 14 days. Upon release from quarantine, mice were housed individually for 7 days to reduce stress prior to calorimetry. Mice were then placed into an indirect calorimetry system (Promethion, Sable Systems, Las Vegas, NV) to measure whole body energy expenditure/metabolism, and voluntary movement/activity (see **Figure 2A** for a graphic summary of the procedure). Mice were acclimated for 3-4 days in the system prior to study data collection. After the acclimation period, data were collected and analyzed with Macro Interpreter (V24.1.0) in 15-minute and 1-hour increments using a One Click Macro provided by Sable Systems (V.2.53.2-slice15mins and V.2.53.2-slice1hr, respectively). Measurements were obtained from 28 individuals aged 3-4 months spread over two experimental cohorts. Cohort 1 consisted of 16 individuals (4 wild-type females, 4 R263Q females, 4 wild-type males and 4 R263Q males) measured for 17 full days, and Cohort 2 comprised 12 individuals (same as Cohort 1 but no R263Q females included) measured for 9 full days. All downstream data processing, analysis and plotting steps were carried out in RStudio (version 4.4.0).

**Figure 2.**
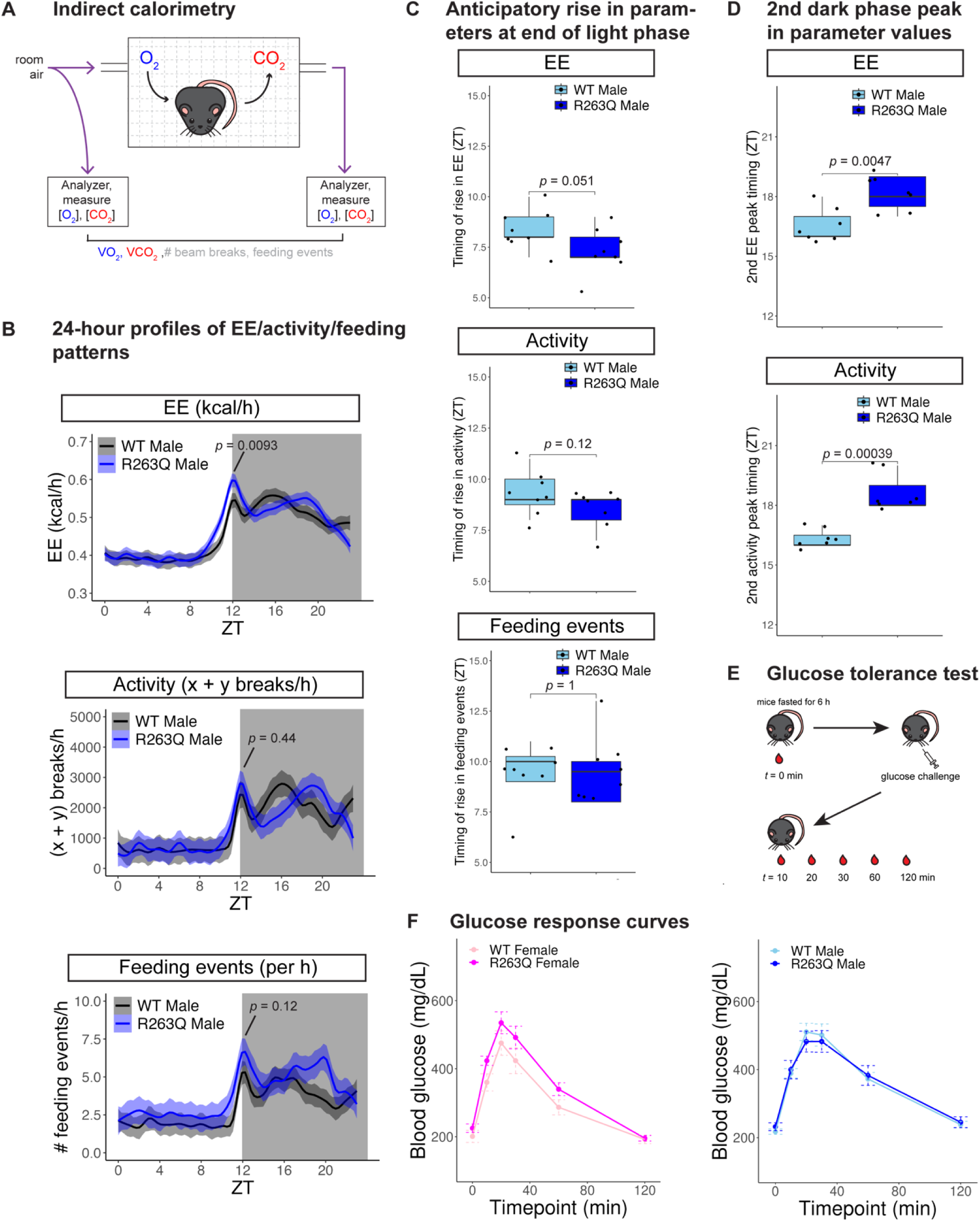
CRY1 R263Q males show delays in 24-hour EE, activity and feeding patterns in the dark phase, but no defects in glucose metabolism in either sex. 2A. Graphic of metabolic phenotyping using indirect calorimetry (IC). Each experimental subject is housed individually in its own metabolic cage. O_2_ and CO_2_ concentrations of air entering the cage and air exiting the cage are measured by analyzers and used to calculate the volume of O_2_ inspired and volume of CO_2_ expired. These values can then be used to infer the resting energy expenditure (EE) of the animal. Each cage is also equipped with a grid of infrared beams (shown in dashed grey lines) in the x-, y– and z-directions. The number of beam breaks is recorded and can be used to infer the locomotor activity (sum of breaks in the x– and y-directions) of the individual. Contact events with the food hopper are also recorded as feeding events. 2B. 24-hour profiles of energy expenditure (EE), activity and feeding events as a function of zeitgeber time (ZT). Values for each metabolic parameter were averaged over the experiment duration to produce a single hourly value for each individual. Solid lines represent means of hourly values computed for all individuals in an experimental group and shaded areas indicate 95% confidence intervals around the means. Wild-type males are represented by black lines with grey shading. R263Q males are represented by blue lines with pale blue shading. The grey rectangle spanning the whole y-axis and going from ZT12-ZT24 represents the dark phase. Top: energy expenditure (EE); middle: activity; bottom: feeding events). *p*-values are obtained from Student’s *t*-test and indicate the comparison between the wild-type and R263Q values recorded at ZT12. *n* = 8 individuals each for wild-type males and R263Q males. 2C. Boxplots comparing the timings (measured in ZT) of the anticipatory rise in EE (top), activity (middle) and feeding events (bottom) at the end of the light phase. *p*-values are obtained from Student’s *t*-test. *n* = 8 individuals each for wild-type males and R263Q males. 2D. Boxplots comparing the timings (measured in ZT) of the second peak in EE (top) and activity (bottom) during the dark phase. *p*-values are obtained from Student’s *t*-test. *n* = 7 individuals each for wild-type males and R263Q males; one individual from each group was left out because identifying the timing of the second peak was not possible for these individuals. 2E. Graphic summary of the glucose tolerance test (GTT) procedure. Mice are fasted for 6 hours, and then fasting blood glucose measurements (t = 0 min) are obtained by pricking the tail vein. Glucose is then administered at a dose of 2 mg/g of body weight by intraperitoneal injection. Blood glucose measurements are then obtained at 10, 20, 30, 60 and 120 minutes following the glucose challenge and then used to plot a glucose response curve for each individual. 2F. Glucose response curves showing blood glucose levels (in mg/dL) as a function of time elapsed since glucose injection. Left: females; right: males. Points represent mean glucose levels for each group at a given time point, error bars represent standard error. *n* = 9 for wild-type females (shown in pink), 7 for R263Q females (shown in magenta), 9 for wild-type males (shown in sky blue) and 7 for R263Q males (shown in blue).

### Glucose tolerance testing

Mice housed at SIMR and aged 2 months were fasted for about 6 hours from 6:30 am to 12:30 pm to acquire fasting blood glucose measurements. Before the start of the experiment, each mouse was weighed and then tail veins were pricked to draw blood that was used to measure fasting blood glucose levels using an Alpha Trak 3 glucometer and test strips. Mice were then injected intraperitoneally with 2 mg/g of body weight of glucose (A24940-01, Gibco) administered as 10 μL per g of body weight of a 20% w/v glucose solution. Postprandial blood glucose levels were then measured at 10, 20, 30, 60 and 120 minutes post glucose injection. Experiments were done in 3 independent cohorts, and then pooled for plotting and analysis, with individuals from each experimental group represented in each cohort. A graphic summary of the GTT procedure is provided in **Figure 2E**.

### RNA extraction and sequencing

Mice housed at SIMR and aged between 3-4 months were euthanized with CO_2_ followed by cervical dislocation. Brains and livers were harvested from each mouse and washed in 1X PBS. The left half of each brain, and fragments of the largest lobe of each liver were immediately homogenized in TRIzol reagent (15596026, ThermoFisher Scientific) using a tissue homogenizer (Z742682, Sigma) with 1.5 mm zirconium beads. Tissue harvesting was done at two zeitgeber times – between ZT12 and ZT13 (onset of the dark phase) and between ZT22 and ZT23 (onset of the light phase). RNA was extracted following the manufacturer’s instructions. The extracted RNA samples were treated with DNase I (M0303S, New England Biolabs) and then purified using the NEB Monarch RNA Cleanup Kit (T2050, New England Biolabs) following the manufacturer’s instructions.

mRNAseq libraries were generated from 500 ng of high-quality total RNA, as assessed using the Bioanalyzer (Agilent), according to the manufacturer’s directions using a 5-fold dilution of the universal adaptor and 9 cycles of PCR per the respective masses with the NEBNext Ultra II Directional RNA Library Prep Kit for Illumina (NEB, E7760L), the NEBNext Poly(A) mRNA Magnetic Isolation Module (NEB, E7490L), and the NEBNext Multiplex Oligos for Illumina (96 Unique Dual Index Primer Pairs) (NEB, E6440S) and purified using the SPRIselect bead-based reagent (Beckman Coulter, B23318). The resulting short fragment libraries were checked for quality and quantity using the Bioanalyzer (Agilent) and Qubit Flex Fluorometer (Life Technologies). Equal molar libraries were pooled, quantified, and converted to process on the Singular Genomics G4 with the SG Library Compatibility Kit, following the “Adapting Libraries for the G4 – Retaining Original Indices” protocol. The converted pools were sequenced on F2 flow cells (700101) on the G4 instrument with the PP1 and PP2 custom index primers included in the SG Library Compatibility Kit (700141), using Instrument Control Software 23.08.1-1 with the following read lengths: 8 bp Index1, 50 bp Read1, 8 bp Index2, and 50 bp Read2. Following sequencing, sgdemux 1.2.0 was run to demultiplex reads for all libraries and generate FASTQ files.

### RNA-seq data analysis

Samples from each ZT were collected, sequenced and analyzed separately. Raw reads were demultiplexed into FASTQ files using Illumina’s bcl-convert (version 3.10.5) before being aligned to UCSC genome GRCm39 with STAR aligner (version 2.7.10b), using Ens_110 gene models. TPM values were generated using RSEM (version v1.3.1). The STAR read counts table was used for differential gene expression analysis using edgeR package (version 3.42.4) in RStudio (R version 4.3.1). Lowly expressed genes were filtered out with filterByExpr’s default settings followed by a trimmed mean of M-values (TMM) normalization. Differential gene expression comparisons were performed between genotypes (wild-type and R263Q) but within the same sex, tissue type and ZT. Genes showing absolute values of log2 fold change > 1 (or absolute values of fold change > 2) and adjusted *p*-values less than 0.05 were identified as differentially expressed. Up– and down-regulated gene lists from each comparison were used as input for Gene Ontology (GO term) pathway enrichment analysis. GO term enrichment was completed using TERMS2GO, an in-house R Shiny app (versions R 4.3.1, shiny 1.7.5). Significant gene ontology terms were identified using clusterProfiler’s enrichGO function (version 4.8.2) with AnnotationHub’s species database (version 3.8.0). GO Terms with adjusted p-values less than 0.05 were considered significant. Figures were generated using ggplot2 (version 3.4.3) and plotly (4.10.2).

### Statistical analyses

For indirect calorimetry and glucose tolerance test analyses, Student’s *t*-test was used to compare wild-type and R263Q samples within sexes. For multiple comparisons (wild-type female versus R263Q female and wild-type male versus R263Q male), the Bonferroni correction was used for adjusting the *p*-values rather than analysis of variance (ANOVA), since ANOVA adjusts for all possible comparisons, whereas we were interested *a priori* in evaluating only the two fixed comparisons mentioned above.

## RESULTS

### A mouse model for the repeatedly evolved CRY1 R263Q mutation

A mutation was recently reported in the gene *Cryptochrome-1* (*Cry1*) to cause an amino acid substitution from a positively charged arginine to an uncharged polar glutamine in a protein domain that is otherwise highly conserved across metazoans (Moran et al., 2022) (**Figure 1A**). Moreover, this mutation has been detected in several species of cave-dwelling fish, namely, *Astyanax mexicanus*, *Sinocyclocheilus spp.*, *Phreatichthys andruzzii*, and in two burrowing mammals, the naked mole-rat *Heterocephalus glaber*, and the degu *Octodon degus*. Therefore, the CRY1 R263Q mutation has arisen independently at least 5 times during animal evolution (**Figure 1B**). Given that this specific mutation has repeatedly occurred in a highly conserved gene, in species that share similar environmental pressures, we hypothesize that this mutation does not result simply in a loss of function. Rather, it might target a specific function of this protein.

To test this hypothesis and investigate the function of the CRY1 R263Q mutation, we used CRISPR/Cas9 to introduce the underlying G788A nucleotide substitution at the endogenous *Cry1* locus of the mouse genome (**Figure S1**, also see Materials and Methods). We performed RNA sequencing (RNA-seq) using brain and liver tissues collected at ZT 12-13 and confirmed that *Cry1* is transcribed at comparable levels in these organs between wild-type and R263Q individuals (**Figure 1C**, left). We then performed western blotting with brain and liver lysates from the same mice using CRY1 antibodies to confirm comparable levels of CRY1 protein expression between wild-type and R263Q individuals (**Figure 1C**, right). Consequently, we conclude the CRY1 R263Q mutation does not impact the normal transcription and translation of the *Cry1* gene i.e. it does not phenocopy a CRY1 loss-of-function.

### CRY1 R263Q mice have delayed energy expenditure and locomotor activity patterns in the dark phase relative to wild-type mice

Since CRY1 is a key regulator of mammalian circadian rhythms and metabolism, we screened the CRY1 R263Q mice and wild-type controls for altered behavioral and metabolic parameters using indirect calorimetry (IC, see Materials and Methods and **Figure 2A**). We collected O_2_ and CO_2_ measurements (mL/min) to infer energy expenditure (EE), as well as locomotor activity (or simply activity) patterns measured as infrared beam breaks in the x– and y-directions, and feeding events (see Materials and Methods for details). We observed no significant changes to gross EE, locomotor activity or feeding frequency (number of feeding events) in the R263Q mice over a full day or during specific photoperiods (**Figure S2**). To refine the temporal resolution of our analyses, we then examined EE patterns hourly over the course of a 24-hour day. We computed the mean values for hourly EE across all the experiment days for each mouse and visualized the values as a function of zeitgeber time (ZT) (**Figure 2B**, top panel). We observed altered EE patterns in the R263Q mutants relative to wild-type controls. For males, during ZT 0-9, the EE remained relatively constant and very similar between wild-type and R263Q (∼0.4 kcal/h). At around ZT9, the EE rose rapidly peaking at ZT12, with R263Q individuals reaching a slightly higher peak compared to the wild-type (**Figure 2B** top panel, *p* = 0.0093). The timing of this anticipatory rise was slightly sooner for the R263Q individuals than the wild-type (**Figure 2C** top panel, *p* = 0.051). Wild-type mice showed a dip at ZT13, a second peak at ZT16 (∼0.6 kcal/h), and then a steady decline until ZT20 before rising again. R263Q individuals exhibited a similar pattern but with significant delay in the timing of the second peak at ZT18-19 (**Figure 2D** top panel, *p* = 0.0047). Females showed similar trends, but the differences between female genotypes did not reach statistical significance (**Figure S3**), except for the timing of anticipatory rise in EE at the end of the light phase. Female R263Q mice showed this increase sooner than wild-type females (**Figure S3B** top panel, *p* = 0.016).

To further investigate the delayed EE patterns in R263Q individuals, we compared 24-hour profiles of activity between wild-type and R263Q males (**Figure 2B**, middle panel). The activity patterns mirrored the 24-hour EE patterns. Both wild-type and R263Q males showed baseline activity (∼500-1000 breaks/h) from ZT0 to ZT10, then increased sharply up to ZT12, without any difference in the timing of this increase (**Figure 2C** middle panel, *p* = 0.12) or in the peak values attained (**Figure 2B** middle panel, *p* = 0.44). Wild-type males showed a dip in activity at ZT13, then a second peak at ZT16 dropping steadily until ZT20 before rising again. R263Q males showed a significantly delayed second peak in activity after ZT12, reminiscent of the EE patterns (**Figure 2D** bottom panel, *p* = 0.00039). The females showed similar patterns as the males, but without statistically significant differences between the genotypes (**Figure S3**), except for the anticipatory increase in activity at the end of the light phase that bordered on statistical significance (**Figure S3B** middle panel, *p* = 0.06).

Next, we examined 24-hour profiles of feeding events. Similar to the EE and activity profiles, wild-type male feeding events remained constant up to ZT11, then rose steeply to peak at ZT12 (∼5 events/h) (**Figure 2B**, bottom panel). After a dip at ZT13, frequencies peaked again at ZT16-17, declined until ZT20, then increased again. The R263Q males mirrored the wild-type from ZT0-12. R263Q mutants, too, increased feeding events starting at ZT9 like the wild-type (**Figure 2C** bottom panel, *p* = 1) and peaked at ZT12 with feeding frequency similar to the wild-type (**Figure 2B** bottom panel, *p* = 0.12). Post-ZT12, R263Q males showed a dip to ∼3 events/h at ZT13, but then increased to a second peak of 7 events/h at ZT20, before dropping until ZT23. R263Q females showed a peak at ZT12 but then a decline up to ZT23, while the wild-type female profile resembled that of the wild-type males (**Figure S3A** bottom panel). No other differences were found between the female genotypes (**Figure S3**). We also compared respiratory exchange ratios (RER) between male and female genotypes and did not detect any differences either in the photoperiod average analysis (**Figure S4A**) or in the 24-hour profiles (**Figure S4B**).

Mutating or depleting mouse CRY1 has been previously reported to cause hyperglycemia and inability to regulate blood glucose levels following a glucose challenge (Lamia et al., 2011; Okano, 2018). This phenotype is present in Mexican cavefish that naturally harbor the R263Q mutation, raising the possibility that this mutation disrupts glucose tolerance (Riddle et al., 2018). To test this hypothesis, we compared R263Q and control mice using a standard glucose tolerance test (see Materials and Methods and **Figure 2E**). We found no significant differences in fasting blood glucose levels between wild-type and R263Q individuals of either sex (**Figure S4C**). After a glucose challenge, all groups showed the expected increase in blood glucose levels in the first 30 minutes and were able to tolerate glucose and regulate blood glucose back to fasting levels within 2 hours of the glucose challenge (**Figure 2F**). These data therefore do not provide any evidence of insulin resistance or impaired glucose homeostasis capacity in the R263Q mice. In summary, male mice carrying the CRY1 R263Q mutation showed a delay in their energy expenditure, activity and feeding patterns in the dark phase. R263Q females show similar trends that do not reach statistical significance. Delayed nocturnal feeding behaviors and locomotor activity patterns in males could reasonably explain the delayed EE patterns. Finally, the R263Q mice do not have deficits in glucose metabolism as assayed by a standard glucose tolerance test.

### CRY1 R263Q mice show altered expression of circadian and metabolic genes in major organs

We next sought to examine the effects (if any) of the CRY1 R263Q mutation on gene expression, to see if it could explain the altered behavioral patterns recorded by indirect calorimetry. Since we were assaying for both circadian and metabolic phenotypes, we selected the brain and liver to perform RNA-seq and search for gene expression differences between wild-type and R263Q mice. The brain contains the suprachiasmatic nucleus (SCN) which is the central circadian pacemaker that can be reset by light (reviewed in (Takahashi, 2017)). The liver, in addition to regulating metabolism, also has a tissue-level peripheral clock that is responsive to non-photic entrainment cues such as feeding (Manella et al., 2021). Moreover, since circadian and metabolic gene expression profiles can vary depending on the time of day and can be set by external cues, we sampled brain and liver tissue at two different zeitgeber times – onset of the dark phase (ZT 12-13) and onset of the light phase (ZT 22-23). The samples collected are shown in **Figure S5A**.

Correlation analyses (**Figure S5B-C**) showed that for each ZT examined, samples clustered by tissue type. The liver samples clustered largely by sex, but there was no evident clustering by genotype within the sexes. The brain samples, on the other hand, did not show a clear clustering either by sex or by genotype. Regardless, we kept the sexes separate throughout the analyses in keeping with the sex differences observed in the indirect calorimetry experiments. The correlation analyses suggest that while sex could have an impact on gene expression, the impact of the genotype on the overall transcriptional profile is mild. Nonetheless, we did find some noteworthy changes to the transcriptome across the comparisons of interest. The most pronounced transcriptional response was in the tissues of the R263Q males relative to wild-type males, specifically at the onset of the dark phase (**Figure 3A**). This is consistent with R263Q males showing a significant behavioral change relative to the wild-type in the dark phase as recorded by indirect calorimetry (**Figure 2**). Some metabolic genes were upregulated in the brains of R263Q males, such as *glutathione peroxidase 3* (*Gpx3*) which is involved in protection against oxidative damage to cells (reviewed in (Zhang et al., 2024)), and *CART prepropeptide* (*Cartpt*), which codes for several peptides that regulate energy balance, appetite and body weight (Lau and Herzog, 2014). Genes downregulated in the brains of R263Q males were enriched mostly in cell division functions (**Figure 3A** bottom left panel). Other downregulated genes also code for DNA– and RNA-binding proteins such as *Srek1*, *Thoc2*, *Luc7l2*, *Pnisr*, *Prpf39* and a variety of zinc finger proteins. In the livers of R263Q males, a core circadian gene, *Arntl*/*Bmal1*, was upregulated (**Figure 3A**, top right panel). The BMAL1 protein is also a major regulator of metabolic processes like gluconeogenesis and lipogenesis (Hatanaka et al., 2010; Shostak et al., 2013). The arginine vasopressin receptor *Avpr1a,* which contributes to rhythmic behavior in mice, was also upregulated (Li et al., 2009). Genes downregulated in R263Q livers were enriched for several metabolic and rhythmic processes including circadian rhythms (**Figure 3A**, bottom right panel). Major circadian genes that were downregulated in the R263Q male liver included *Nr1d2*, *Dbp*, *Noct* and *Ciart* (**Figure 3A**, top right panel). The *Nr1d2* and *Dbp* genes function in the molecular circadian oscillator and can affect the period and phase of the molecular clock (Takahashi, 2017). *Noct* in humans is under circadian control and regulates metabolism by dephosphorylating NADPH (Estrella et al., 2019). The *Ciart* gene is predicted to be a regulator of circadian gene expression (Annayev et al., 2014). Several metabolic genes were also downregulated, including regulators of lipid and steroid metabolism such as coenzyme Q10B (*Coq10b*), lipase G (*Lipg*), retinol dehydrogenase 9 (*Rdh9*) and 16 (*Rdh16*), glycerophosphocholine phosphodiesterase 1 (*Gpcpd1*), uridine phosphorylase 2 (*Upp2*), and phosphoenolpyruvate carboxykinase 1 (*Pck1*), the latter being a key regulator of gluconeogenesis (Hanson and Garber, 1972). Females showed almost no differentially expressed genes between genotypes at the onset of the dark phase (**Figure S6A**, left panels).

**Figure 3.**
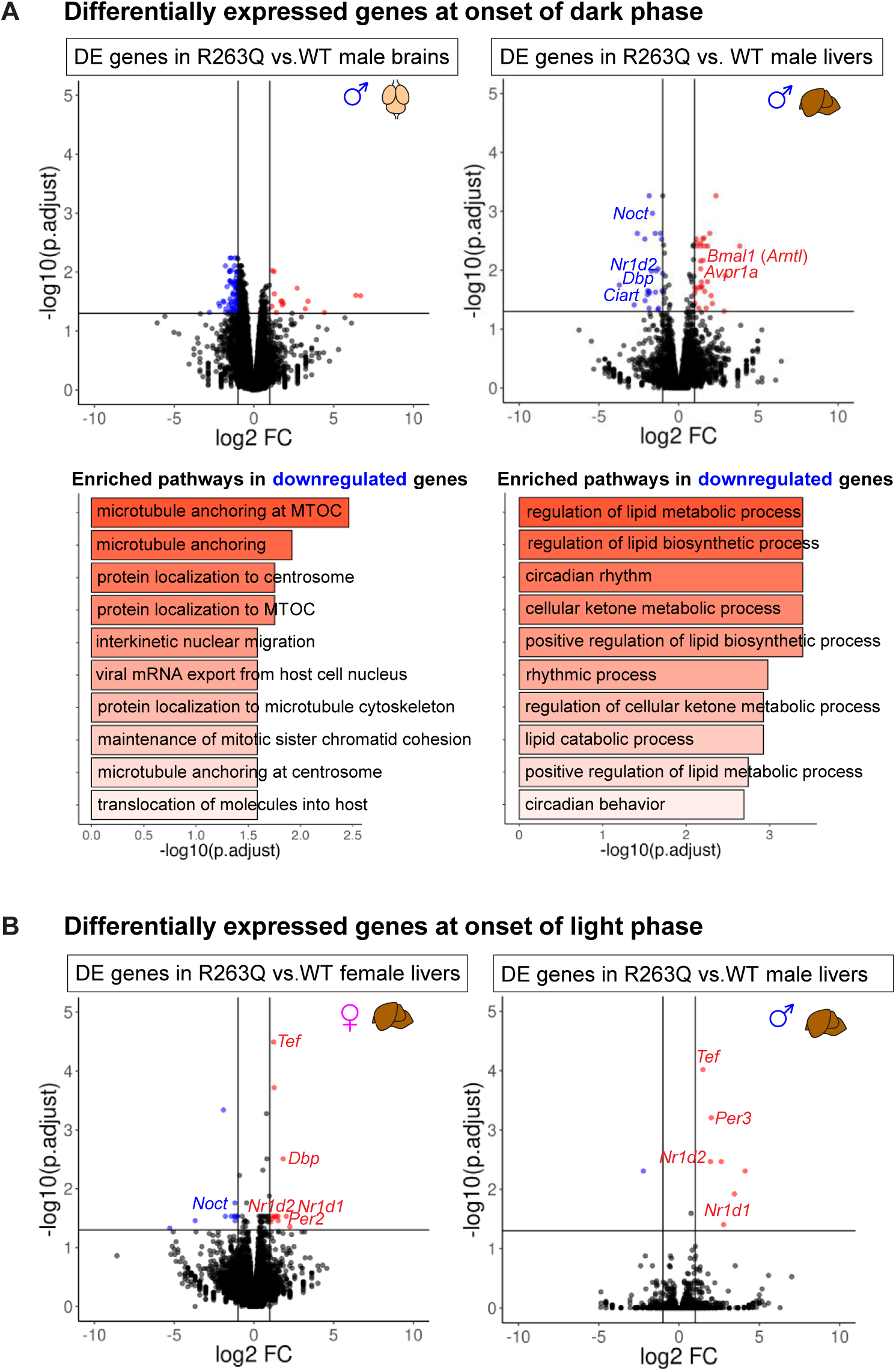
RNA-seq shows altered expression of canonical circadian genes in tissues of CRY1 R263Q mice. 3A. Analysis of genes differentially expressed in brain and liver tissues of male mice (R263Q vs. wild-type) at the onset of the dark phase. Top: volcano plots showing –log10 of adjusted *p*-value versus log2 fold change for genes expressed in the brains (top left) and livers (top right) of R263Q males relative to the wild-type. Bottom: Gene Ontology (GO term) enrichment analysis of biological pathways overrepresented in genes that are downregulated in the brains (bottom left) and livers (bottom right) of R263Q male mice relative to the wild-type. 3B. Volcano plots showing –log10 of adjusted *p*-value versus log2 fold change for genes expressed in the livers of R263Q individuals relative to the wild-type at the onset of the light phase. Left: females; right: males. For all volcano plots shown, vertical lines indicate cut-off for fold change (±2) for a gene to be considered differentially expressed, and the horizontal line indicates the cut-off for adjusted *p*-value (0.05) for a gene to be considered differentially expressed. Upregulated genes are represented by red points and downregulated genes by blue points.

At the onset of the light phase, a modest transcriptional response was observed in the livers of R263Q mice relative to the wild-type (**Figure 3B**), but almost none in the brains (**Figure S6A**, right panels). For both sexes, genes upregulated in the mutant livers included canonical circadian genes such as *Nr1d1/2*, *Tef*, *Dbp* and *Per2/3*, all of which are involved directly in the molecular circadian oscillator. In addition to circadian processes, genes upregulated in the livers of R263Q females were also enriched for various metabolic processes (**Figure S6B**). Upregulated metabolic genes included acyl-CoA thioesterase 1 (*Acot1*) that is involved in fatty acid metabolic processes (Franklin et al., 2017), and cytochrome P450 family 26 subfamily B member 1 (*Cyp26b1*), a monooxygenase enzyme whose expression is regulated by retinoic acid (Loudig et al., 2000). Genes downregulated in R263Q female livers were mainly enriched for metabolic processes such as detoxification and response to toxic substances, and cellular response to metal ions (**Figure S6B**). These genes include metallothionein 1/2 (*Mt1*/*2*) and cytochrome P450 family 2 subfamily a polypeptide 5 (*Cyp2a5*). In summary, R263Q animals show aberrant circadian and metabolic gene expression profiles in their brain and liver tissues, with circadian gene dysregulation restricted to the liver tissues. This aligns with the observed behavioral and metabolic changes exhibited by these animals.

## DISCUSSION

In this study, we investigated the transcriptomic, behavioral, metabolic and physiological consequences of a repeatedly evolved mutation (R263Q) in the circadian protein Cryptochrome-1, or CRY1. This mutation resides in a highly conserved domain of the protein and is seen to recur specifically in animal lineages that have adapted to low-light environments like caves and subterranean burrows. We developed a mouse model that is homozygous for the CRY1 R263Q mutation and carried out an in-depth metabolic phenotyping to uncover the potential effects of this mutation. We found a change in 24-hour EE profiles in CRY1 R263Q mutants that are likely caused by similar changes in locomotor activity and feeding behavior. Finally, RNA sequencing revealed disruption in expression patterns of several canonical circadian and metabolic genes. Through this study, we have demonstrated the power of a single nucleotide substitution to orchestrate change not only at the level of gene expression, but also at the level of the whole organism.

### Does the CRY1 R263Q mutation phenocopy wild-type CRY1, CRY1 loss of function, gain of function or neofunctionalization?

The evolutionary convergence of the CRY1 R263Q mutation suggests it that does not represent a complete loss of function and might instead be selected for. Here, we provide multiple lines of evidence combining our findings with the existing literature to argue that CRY1 R263Q retains much of its function, indicating a partial loss of function or neofunctionalization of the CRY1 protein.

Under conditions of constant darkness, *Cry1*^−/−^ mice show a shortened free-running period relative to wild-type mice (Griffin et al., 1999; Kume et al., 1999; van der Horst et al., 1999), which is also reflected in the molecular rhythms of the *Per1* gene (Griffin et al., 1999). Gul *et al*. (Gul et al., 2020) found that substituting CRY1 Arg293 with His (CRY1 R293H) leads to a shortened molecular rhythm, suggesting that the CRY1 R293H functions as a hypomorph. There are also other mutations in the literature that suggest a CRY1 hypermorphic phenotype. Replacing CRY1 Cys414 with Ala (CRY1 C414A) leads to a longer free-running period in darkness (Okano et al., 2009), with reduced affinity for its binding partner PER2 (Schmalen et al., 2014). Another variation of CRY1, consisting of an in-frame deletion of 24 amino acids coded in exon 11, shows a similar period-lengthening phenotype and is implicated in familial delayed sleep phase disorder (DSPD) (Patke et al., 2017). The CRY1 R263Q mutation which we have characterized in this study does not clearly phenocopy either the full loss of function, the hypomorph or the hypermorph phenotype. However, it is important to note that our indirect calorimetry and activity measurement experiments for the CRY1 R263Q mutation were carried out in a standard light/dark cycle. Since light can mask a malfunctioning biological oscillator by acting as a zeitgeber to provide an extrinsic rhythmic cue (van der Horst et al., 1999), we cannot make powerful statements about possible changes to circadian period lengths of CRY1 R263Q mice under dark conditions. However, there are still several noteworthy observations that we made for the CRY1 R263Q mutation. One, the CRY1 R263Q mice showed a delay in locomotor activity patterns and consequently the energy expenditure patterns in the dark phase. It is therefore tempting to speculate that this delay would manifest as a lengthened circadian period under total darkness. This could be linked to reduced PER2 affinity as reported for the CRY1 C414A mutation (Schmalen et al., 2014), or due to a perturbation to the balance between competitive FAD and ubiquitin ligase binding to regulate CRY1 stability (Busino et al., 2007; Hirano et al., 2017; Hirano et al., 2013), as a consequence of the proximity of the mutation site to the FAD/FBXL3-binding site of CRY1 (Moran et al., 2022). Further experiments to determine binding partners of CRY1 R263Q, coupled with a behavioral phenotyping in total darkness, would be required to fully elucidate the mechanism by which the CRY1 R263Q mutation induces a phase shift in activity and energy expenditure patterns. Additionally, phenotyping behavior in darkness will also allow us to tease apart the role of an extrinsic cue such as light, and the intrinsic molecular oscillator, in causing the observed delay phenotype.

Two, we performed a glucose tolerance test on our CRY1 R263Q mice and detected no signs of hyperglycemia or glucose intolerance, consistent with our conclusion that the CRY1 R263Q mutation does not mimic a canonical loss of function. A final crucial point to note is that even the full *Cry1*^−/−^ deletion does not show any discernible alterations to circadian wheel-running patterns in a light/dark cycle, ostensibly due to the masking effect of light as described above (van der Horst et al., 1999). The CRY1 R263Q mutation, however, shows an activity pattern phenotype even under a light/dark cycle. It seems therefore that the effect of this mutation is strong enough to not be masked by light. To the best of our knowledge, this is unlike any CRY1 mutation described thus far and supports our hypothesis that the CRY1 R263Q mutation is a neofunctionalization.

### The CRY1 R263Q mutation contributes to circadian clock dysregulation in the liver

RNA sequencing from major organs shows a clear disruption of circadian gene expression in the CRY1 R263Q mutant strain. First, it is interesting to note that the mutation does not significantly alter the expression of the *Cry1* gene itself. This suggests that the mutation exerts its effect at the level of protein function and modulates the expression of other genes directly or indirectly. This hypothesis can be tested by performing chromatin immunoprecipitation sequencing (ChIP-seq) to identify loci bound differentially by wild-type versus R263Q mutant CRY1 and correlate it with the RNA sequencing results.

Second, the RNA sequencing also revealed circadian and metabolic dysregulation as evidenced by altered expression of various components of the interlocked feedback loops that form the molecular circadian oscillator. However, the nature of the differential expression depends on the sex, the tissue type and the zeitgeber time at which RNA was isolated. Circadian genes are not differentially expressed in either of the female tissues examined at the onset of the dark phase. In males, at the onset of the dark phase, circadian genes are not expressed differentially in the brain but there are differences in the liver. Most of the differentially expressed circadian genes are downregulated in the livers of the R263Q males. At the onset of the light phase, there are just a few genes that are differentially expressed in the female brain, and none in the male brain. The livers of the mutant mice, however, in the case of either sex, show a general upregulation of major circadian genes. There are two major points to note here. The first is that the differential expression of circadian genes was identified only in the liver, with no such differences in the brain, irrespective of sex or zeitgeber time. This points to the possibility that the CRY1 R263Q mutation specifically affects peripheral clocks without necessarily disrupting the central circadian pacemaker in the brain. It is conceivable, therefore, that changes to the gene expression profile of the livers could be responsible for altering the feeding patterns of the animals as seen from the indirect calorimetry phenotyping, which could be partially driving the changes in 24-hour locomotor activity patterns and consequently the energy expenditure. The second point of note is that the R263Q males show a general downregulation of circadian genes in the liver at the onset of the dark phase, and both sexes show a general upregulation of circadian genes in the liver at the onset of the light phase. The behavioral data collected from the indirect calorimetry experiments do not show appreciable differences between the wild-type and the R263Q mice at the onset of the light phase, possibly due to the masking effect of light exposure as discussed above. So, even with the upregulation of circadian genes in the liver at the onset of the light phase, a behavioral phenotype if any could be hidden. A future experiment to address this, as noted previously in this section, would be to perform the indirect calorimetry phenotyping under conditions of constant darkness. At the onset of the dark phase, both sexes and genotypes show an anticipatory increase in feeding bouts/activity/energy expenditure (**Figures 2B**, **S3**), and the differences in these patterns become apparent only in the later stages of the dark phase. However, the downregulation of circadian genes in the liver at the onset of the dark phase could be eventually driving the behavioral phase delays that are observed in the R263Q male mice later in the dark phase. At this zeitgeber time, the downregulation of circadian genes is seen only in male R263Q mouse livers, which is consistent with a stronger behavioral phenotype recorded in R263Q males than in females.

### A dysregulated circadian clock in low-light environments

Cave-dwelling species such as the Mexican cavefish *Astyanax mexicanus* and the Somalian cavefish *Phreatichthys andruzzii* are reported to have dysregulated circadian clocks ((Beale et al., 2013; Cavallari et al., 2011; Frøland Steindal et al., 2018; Mack et al., 2021; Olsen et al., 2023)). Additionally, *A. mexicanus* also shows a suite of metabolic traits that resemble disease phenotypes in the human context, such as insulin resistance and increased fat accumulation (Aspiras et al., 2015; Riddle et al., 2018; Xiong et al., 2018; Xiong et al., 2022). Our work shows how a single mutation in the *Cry1* gene leads to aberrant circadian and metabolic gene expression profiles, and delay in nocturnal behaviors.

It is possible that circadian clock dysregulation in low-light environments could be a consequence of relaxed selection, as has been argued for several cave-associated traits (reviewed in (Swaminathan et al., 2023)). If it was indeed a case of relaxed selection, however, then it is extremely unlikely that the very same mutation would occur independently in multiple lineages. This supports the possibility that the R263Q mutation in CRY1 was selected for due to some adaptive advantage offered by a dysregulated circadian clock. Circadian clock dysregulation has been proposed to be adaptive for *A. mexicanus*, for example, since it can serve to conserve energy in a nutrient-limited environment by eliminating rhythmic metabolism (Moran et al., 2014). Future work will examine the precise mechanisms by which the CRY1 R263Q mutation dysregulates molecular and behavioral rhythms and assess its adaptive value in a low-light environment.

Taken together, we have generated a knock-in mouse harboring a CRY1 mutation that has evolved independently multiple times in fish and mammals. The generation of this mouse line allows for systematic investigation of the functional differences, and ultimately, the adaptive nature, of these changes. Our study represents the first functional characterization of a novel, repeatedly evolved mutation in lineages that inhabit cave and subterranean environments. Our analyses provide evidence for changes in metabolic and circadian regulation at multiple levels of organization, providing the basis for future studies examining the role of CRY1 in the repeated evolution of adaptation to low-light environments. Moreover, while the phenomenon of repeated evolution of similar phenotypic traits in response to similar environmental pressures or constraints is not an unfamiliar one, the underlying genetic bases for these shared phenotypes remain elusive. Often, different genetic pathways involved in producing the same phenotype are modified in different lineages to give rise to a nevertheless convergent phenotype. Our work, however, highlights the possibility that the very same genetic change can underly parallel phenotypic changes. In a broader context, our work supports the notion that shared selective pressures can act by favoring the same genotype repeatedly in multiple lineages to drive the evolution of a shared phenotypic trait.

## ACKNOWLEDGEMENTS

This work was supported by institutional funding, NIH New Innovator Award 1DP2AG071466-01, and NIH grants R24OD030214 and P20GM144269. We would like to thank members of the Rohner lab, particularly Fanning Xia, and faculty members of the Graduate School of the Stowers Institute for Medical Research, Dr. Ariel Bazzini, Dr. Matthew Gibson, Dr. Ron Yu and Dr. SaraH Zanders; for regular inputs and feedback on the project. We acknowledge the following members of various technology centers at the Stowers Institute for Medical Research for their valuable services – Heidi Monnin, Marina Thexton and Lana Worthen for animal maintenance and experimental assistance; Madelaine Gogol and Hua Li for general inputs on RNA-seq and statistical analyses; Andrea Moran, Brandon Miller and MaryEllen Kirkman for helping create the CRY1 R263Q line; Sean McGrath and Michael Peterson for generating RNA-seq libraries and sequencing reads; Olga Kenzior, Fengyan Deng and Chongbei Zhao for generating controls to validate the anti-mCRY1 antibodies used in this study; and the Media Prep and Glassware facilities. We also thank Dr. Aziz Sancar and Dr. Christopher Selby (University of North Carolina Chapel Hill, NC, USA) for kindly sharing the pMC1/2SG5 plasmids coding for mCRY1/2 that were used to generate antibody controls. We are also grateful to the Metabolic and Obesity Phenotyping (MORPh) facility at the University of Kansas Medical Center, particularly Dr. John Thyfault, for discussing the data interpretation with us.

---

**Figure S1.**
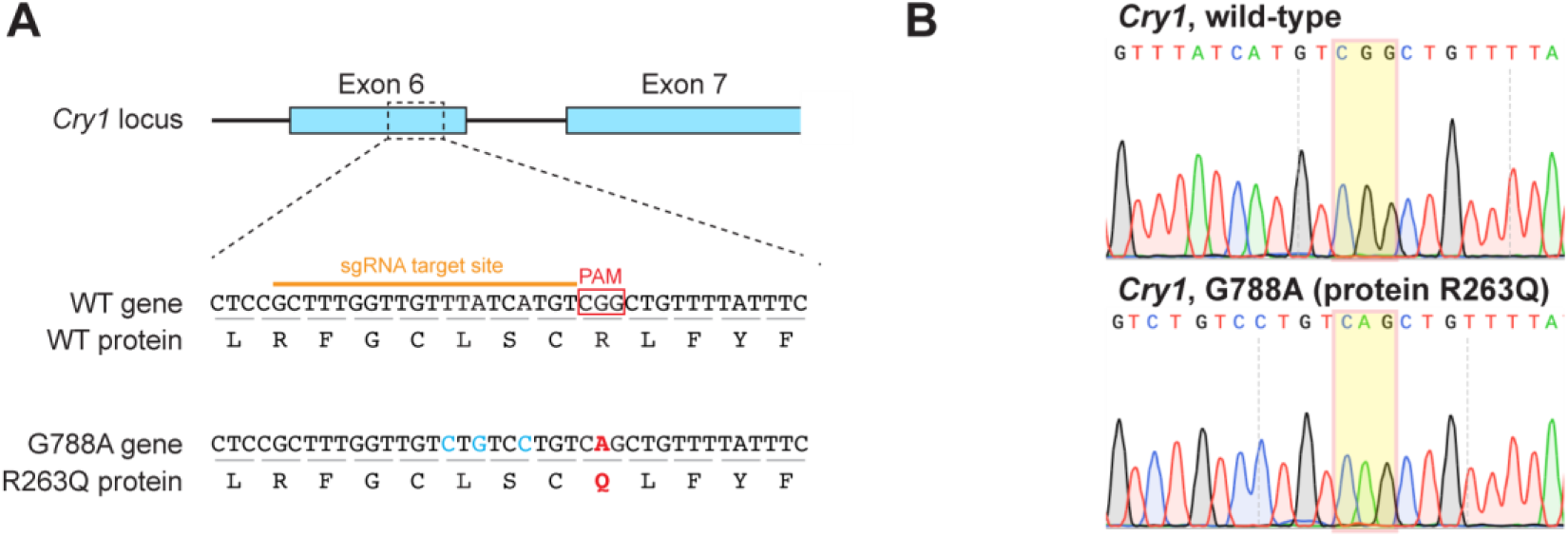
Generation of a mouse model for the CRY1 R263Q mutation using CRISPR/Cas9 technology. S1A. Graphic showing the location of the edits made to the C57BL6/J mouse genome at the *Cry1* locus. The protospacer adjacent motif (PAM) site 5’CGG3’ (red) was incidentally the same codon that was to be edited. The single guide RNA (sgRNA) site is marked in orange. Side-by-side comparisons of the wild-type and G788A mutant coding sequences, and the wild-type and R263Q mutant protein sequences are shown. The mutant sequence contains a single nucleotide change in the *Cry1* coding sequence, G788A, that leads to the intended amino acid substitution R263Q. Additionally, three other silent mutations T778C, A780G and A783C were also introduced to prevent repeated binding of the sgRNA after the desired mutation had been introduced. S1B. Sanger sequencing chromatogram of PCR-amplified region of the G788A mutation site in the *Cry1* locus from a wild-type (top) and a CRY1 R263Q homozygous (bottom) individual. The site of the nonsynonymous G788A mutation is highlighted by yellow boxes. The silent mutations (see Figure S1A) are also present in the CRY1 R263Q chromatogram.

**Figure S2.**
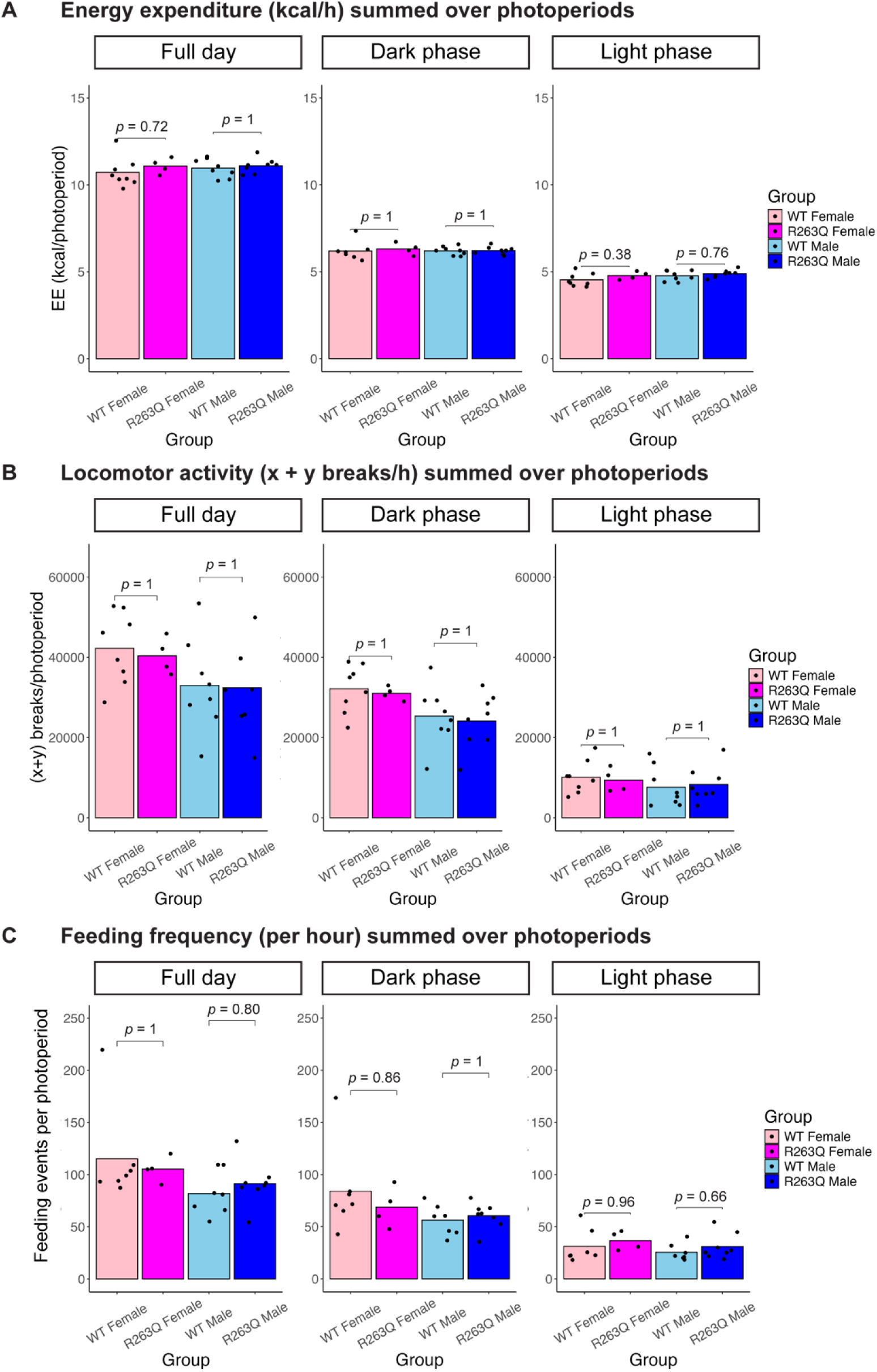
Energy expenditure, locomotor activity and feeding frequencies averaged over a photoperiod do not differ between genotypes. S2A. Energy expenditure (EE) measured in kcal/h summed over three photoperiods (full day, dark phase and light phase) for all four groups of mice assessed. Left: full day; middle: dark phase; right: light phase. S2B. Locomotor activity measured in (x + y) breaks/h summed over three photoperiods (full day, dark phase and light phase) for all four groups of mice assessed. Left: full day; middle: dark phase; right: light phase. S2C. Feeding frequency measured in h^-1^ summed over three photoperiods (full day, dark phase and light phase) for all four groups of mice assessed. Left: full day; middle: dark phase; right: light phase. For all panels, *p*-values are calculated from Student’s *t*-test and adjusted for multiple comparisons using the Bonferroni correction. *n* = 8 for wild-type females, 4 for R263Q females, 8 for wild-type males and 8 for R263Q males.

**Figure S3.**
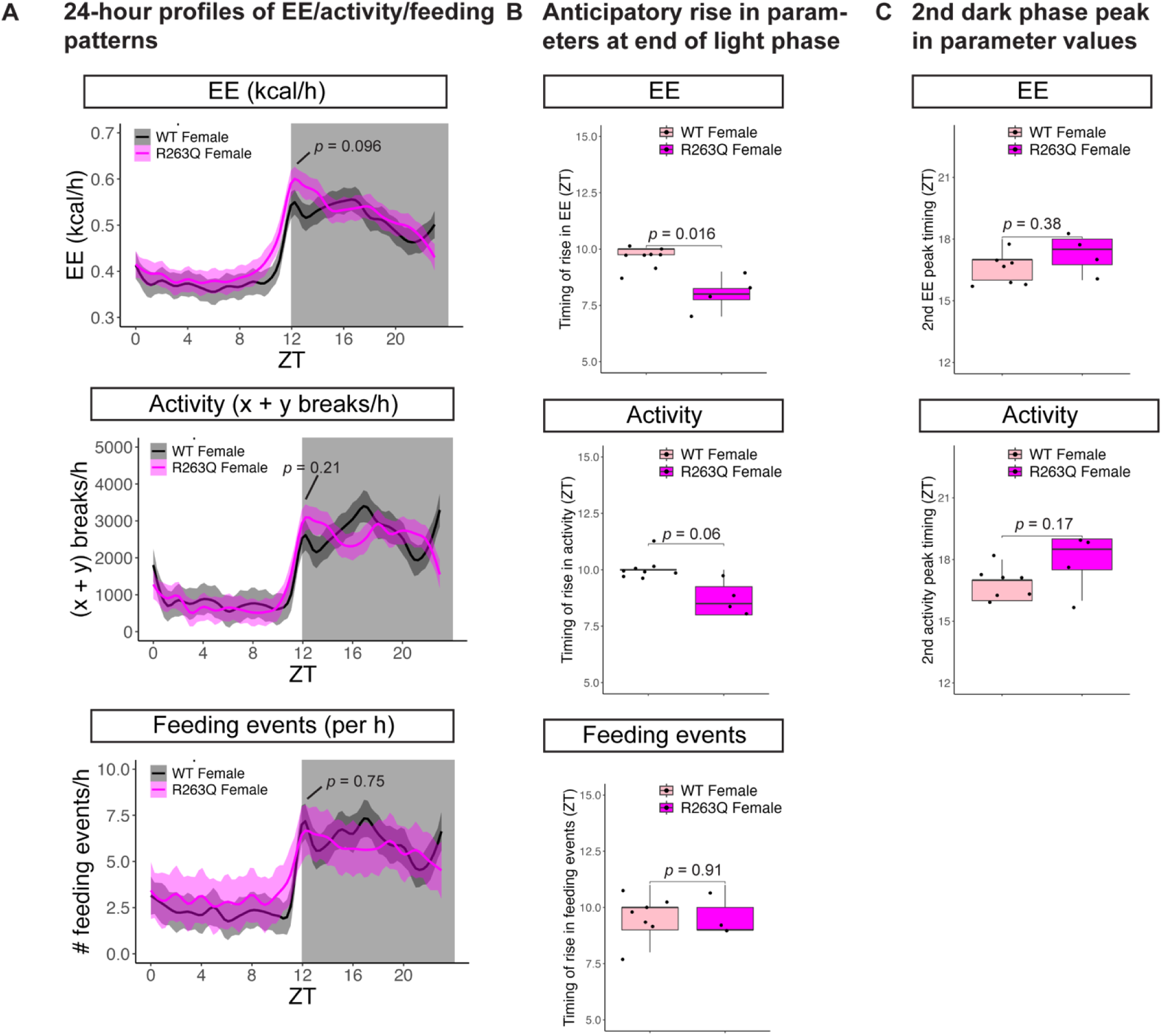
Indirect calorimetry phenotyping for CRY1 R263Q females. S3A. 24-hour profiles of energy expenditure (EE), activity and feeding events as a function of zeitgeber time (ZT). Values for each metabolic parameter were averaged over the experiment duration to produce a single hourly value for each individual. Solid lines represent means of hourly values computed for all individuals in an experimental group and shaded areas indicate 95% confidence intervals around the means. Wild-type females are represented by black lines with grey shading. R263Q females are represented by magenta lines with pale magenta shading. The grey rectangle spanning the whole y-axis and going from ZT12-ZT24 represents the dark phase. Top: energy expenditure (EE); middle: activity; bottom: feeding events). *p*-values are obtained from Student’s *t*-test and indicate the comparison between the wild-type and R263Q values recorded at ZT12. *n* = 8 individuals for wild-type females and 4 for R263Q females. S3B. Boxplots comparing the timings (measured in ZT) of the anticipatory rise in EE (top), activity (middle) and feeding events (bottom) at the end of the light phase. *p*-values are obtained from Student’s *t*-test. *n* = 8 individuals for wild-type females and 4 for R263Q females for anticipatory rise in EE and locomotor activity, and *n* = 7 for wild-type females and 3 for R263Q females for anticipatory rise in feeding events, one individual from each group was left out because identifying the timing of anticipatory rise in feeding events was not possible for these individuals. S3C. Boxplots comparing the timings (measured in ZT) of the second peak in EE (top) and activity (bottom) during the dark phase. *p*-values are obtained from Student’s *t*-test. *n* = 7 individuals for wild-type females and 4 for R263Q females, one wild-type female was left out because identifying the timing of the second peak was not possible for this individual.

**Figure S4.**
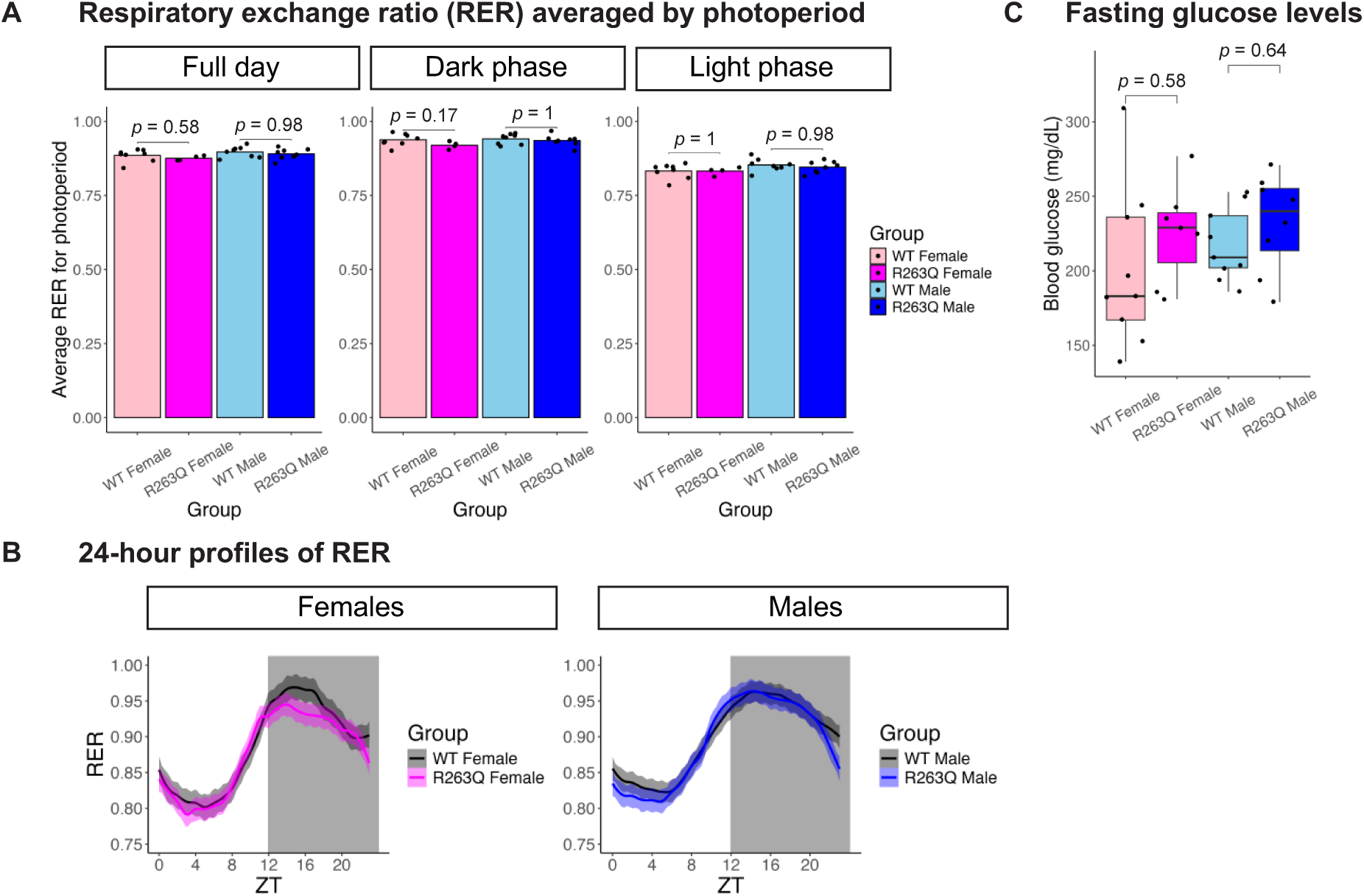
RER and fasting blood glucose levels do not differ between genotypes. S4A. Respiratory exchange ratio (RER) averaged over three photoperiods (full day, dark phase and light phase) for all four groups of mice assessed. Left: full day; middle: dark phase; right: light phase. *p*-values are calculated from Student’s *t*-test and adjusted for multiple comparisons using the Bonferroni correction. *n* = 8 for wild-type females, 4 for R263Q females, 8 for wild-type males and 8 for R263Q males. S4B. 24-hour profiles of RER as a function of zeitgeber time (ZT). RER values were averaged over the experiment duration to produce a single hourly value for each individual. Solid lines represent means of hourly values computed for each individual in an experimental group and shaded areas indicate 95% confidence intervals around the means. Wild-type values are represented by black lines with grey shading. R263Q females are represented by magenta lines with pale magenta shading and R263Q males are represented by blue lines with pale blue shading. The grey rectangle spanning the whole y-axis and going from ZT12-ZT24 represents the dark phase. Left: females (wild-type and R263Q); right: males (wild-type and R263Q). *n* = 8 for wild-type females, 4 for R263Q females, 8 for wild-type males and 8 for R263Q males. S4C. Boxplot showing fasting blood glucose levels for all four groups of mice assessed (wild-type female, R263Q female, wild-type male and R263Q male) as part of the glucose tolerance test shown in Figure 2E-F. *p*-values are calculated from Student’s *t*-test and adjusted for multiple comparisons using the Bonferroni correction. *n* = 9 for wild-type females, 7 for R263Q females, 9 for wild-type males and 7 for R263Q males.

**Figure S5.**
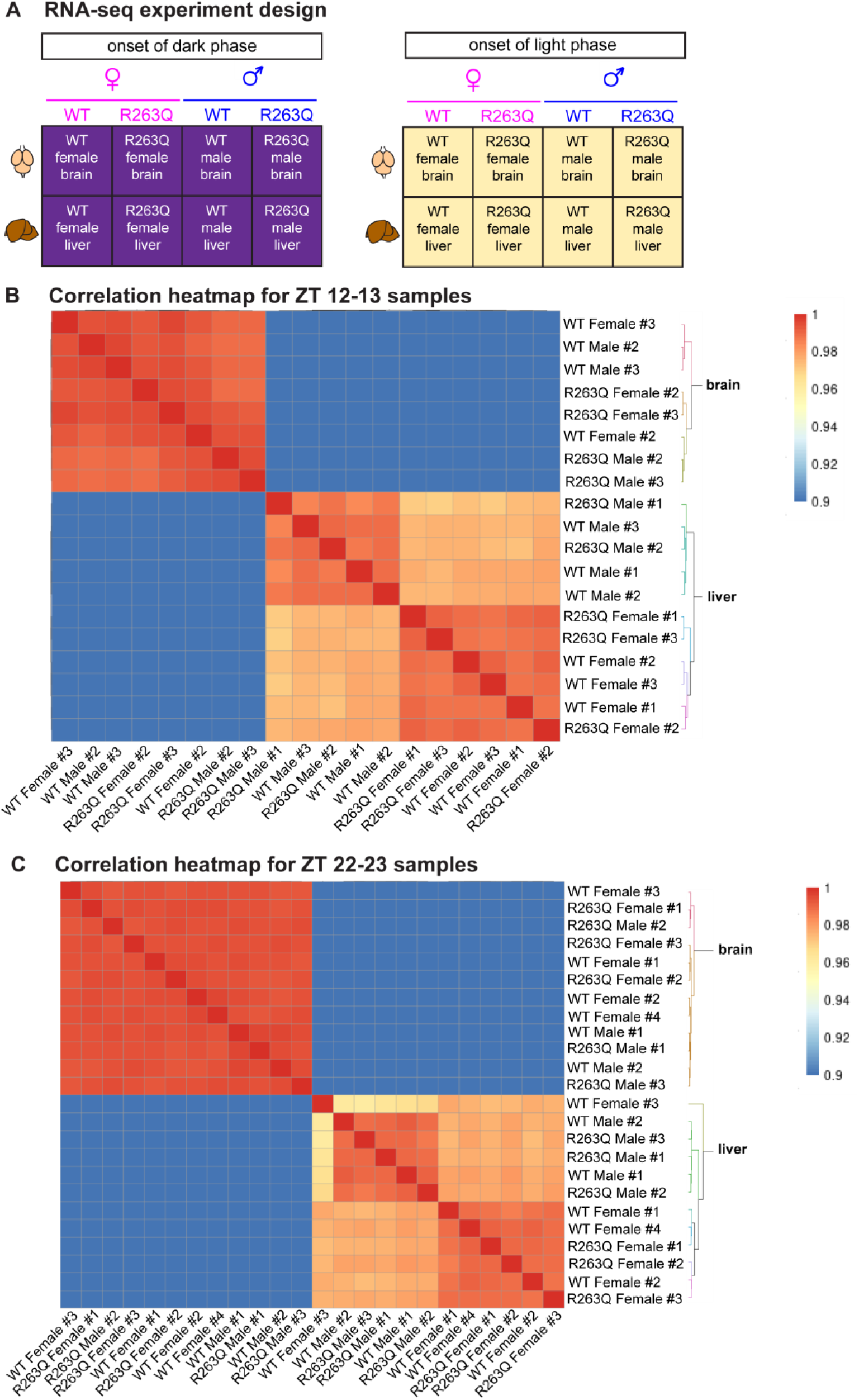
RNA-seq from brain and liver tissues of wild-type and CRY1 R263Q mice at two zeitgeber times. S5A. Experimental design of the RNA-seq experiment showing ZTs, tissues, sexes and genotypes sampled. S5B. Heatmap showing correlation between counts per million (CPM) values for each gene, comparing all samples and replicates collected at the onset of the dark phase (ZT 12-13). *n* = 2 biological replicates for brain samples from each sex/genotype combination, and *3* biological replicates for liver samples from each sex/genotype combination. S5C. Heatmap showing correlation between counts per million (CPM) values for each gene, comparing all samples and replicates collected at the onset of the light phase (ZT 22-23). *n* = 4 biological replicates for each wild-type female tissue type, 2 biological replicates for each wild-type male tissue type, 3 biological replicates for each R263Q female tissue type and 3 biological replicates for each R263Q male tissue type. For both RNA-seq experiments (different ZTs), each biological replicate is an individual. The serial number for each sample label represents the individual from which the sample was derived within that experiment. Color scales for Spearman correlation values are indicated.

**Figure S6.**
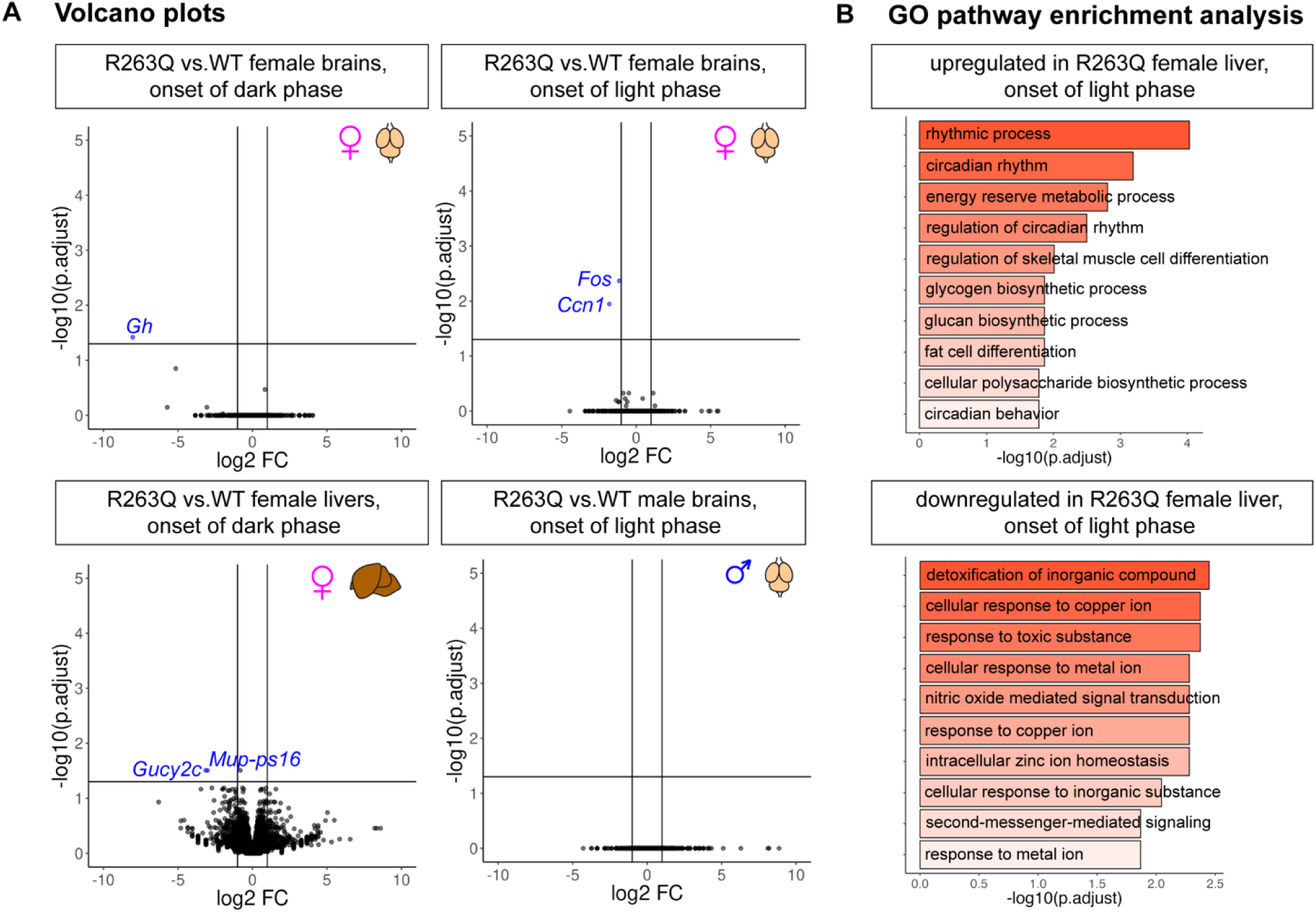
RNA-seq analysis continued. S6A. Volcano plots showing –log10 of adjusted *p*-value versus log2 fold change for expressed genes, for the comparisons not shown in Figure 3. Top left: R263Q vs. wild-type female brains at the onset of the dark phase. Top right: R263Q vs. wild-type female brains at the onset of the light phase. Bottom left: R263Q vs. wild-type female livers at the onset of the dark phase. Bottom right: R263Q vs. wild-type male brains at the onset of the light phase. Vertical lines indicate cut-off for fold change (±2) for a gene to be considered differentially expressed, and the horizontal line indicates the cut-off for adjusted *p*-value (0.05) for a gene to be considered differentially expressed. Upregulated genes are represented by red points and downregulated genes by blue points. S6B. Gene Ontology (GO term) enrichment analysis of biological pathways overrepresented in genes that are up– and down-regulated in livers of R263Q females relative to the wild-type at the onset of the light phase (see Figure 3B for the corresponding volcano plot).

